# A New Evidence of Lubrication Hypothesis on *Nephila pilipes* Spider Silk Spinning

**DOI:** 10.1101/188771

**Authors:** Hsuan-Chen Wu, Shang-Ru Wu, Jen-Chang Yang

**Author notes:** Dr. Jen-Chang Yang will be responsible for all correspondence and correction of the proofs: Graduate Institute of Nanomedicine and Medical Engineering, College of Biomedical Engineering, Taipei Medical University, Taipei, Taiwan 250 Wu-Hsing Street, Taipei 110-52, Taiwan, ROC.

## Abstract

In spite of all the efforts on deciphering such spinning process of spiders, the underlying mechanism currently is yet to be fully revealed. In this research, we designed a novel approach to quantitatively estimate the overall concentration change of spider silk along the progression of liquid-to-solid silk transition from the gland silk. As a prior characterization, we first studied the influence of silking-rate, ranged from 1.5 to 8.0 m/min, on spun fiber diameters as well as fiber strengths. Furthermore, the liquid contents of silk in the sac and the silk fibers leaving the spinneret were investigated by thermogravimetric analysis (TGA) and by estimating the ratio of collected dried silk to the weight loss of spider, respectively. The strength of spun silk fiber showed in the range of 7.5 - 8.5 g/denier; while, the fiber diameter of 0.7 - 1.1 deniers for spun silk first increased then decreased with take-up speed of winder. The results showed that the percentage liquid content of silk stored in the major ampullate sac (50.0 wt%) was lower than that of silk leaving the spinnerets (80.9 - 96.1 wt%), indicating a liquid supplying mechanism might be involved during the spinning process. Thus, a hypothesis of liquid coating on the outer surface of the silk thread served as a lubrication layer to reduce the silking resistance in spinning spigot of spider was proposed. In addition, we speculated the spigot serves as a valve-like regulator that controls not only the fiber diameter but also the lubrication layer. These findings provide understanding in physiological function of the spider spinning process and could further shed some light on future biomimetic development of silk material fabrication.

## Introduction

Studies of spider silk fibers are greatly motivated by their exceptional strength, toughness, biocompatibilities, and potential medical applications (Knight and Vollrath, 2002; Lazaris et al., 2002; Liivak, 1998; Seidel, 1998; Shao and Vollrath, 2002). In spite of significant efforts on mimicking native spider silk fibers, the mechanical performances of the artificially-generated ones, spun from either recombinant or natural silk origin, are often far below the expectation of super strong fibers (Hauptmann et al., 2013; Lazaris et al., 2002; Seidel, 1998; Shao and Vollrath, 2002; Vollrath, 2000; Vollrath and Knight, 1999; Vollrath and Knight, 2001a; Vollrath et al., 2001; Xia et al., 2010; Yao et al., 2002). That is, the source of silk proteins is not the only crucial factor but the processing of the silk materials has also been considered as another indiscernible step towards creating high performance silk fibers (Andersson et al., 2017; Scheibel, 2004; Tokareva et al., 2014; Xia et al., 2010). Spiders harness unique and mild processes, aqueous and adequate temperature, for spinning their silk fibers (Heim et al., 2009; Rammensee et al., 2008). Understanding the silk spinning process of spiders, the naturally born spinners, could provide an essential piece of blueprint towards the fabrication of novel artificially spun materials with superior properties.

Up until now, the spinning process of spider silk within the silk glands is not fully elucidated yet (Koeppel and Holland, 2017). There is a pressing need for more comprehensive understanding regarding the physiological function of a spider’s duct as well as the liquid silk transition within it. The typical ampullate gland consists of a tubular tail portion, a sac-like midpiece and a thin looped spinning duct linking the fluid silk reservoir with the spinneret. The mechanism of silk fiber formation, yet to be completely understood, is generally associated with shear stress induction, pH and ionic gradients, and silk dope concentration change inside the spinning duct (Andersson et al., 2016; Knight and Vollrath, 2001). It is also widely considered that the function of the spinning duct is to extract water out of the liquid silk of a spider (Bell and Peakall, 1969; Davies et al., 2013; Kovoor and Zylberberg, 1972; Tillinghast, 1984; Wilson, 1962; Witt, 1968) during its silking process. Particularly, Vollrath & Knight (Vollrath and Knight, 1999) examined the ultrastructure of spinning duct and indicated that the duct’s wall behaves like a hollow fiber dialysis membrane that removes water from the liquid silk during the spinning. The water-removal role of the spinning duct indeed could beneficially facilitate the silk thread formation, liquid-to-solid silk transition. Subsequently, various mechanisms of liquid crystalline spinning (Vollrath and Knight, 2001a), stress-induced phase separation (Chen et al., 2002; Knight et al., 2000), model of shear-induced conformational change (C. Viney, 1994), and the element composition changes along the spinning duct during the formation of silk threads (Knight and Vollrath, 2001) were all soundly investigated.

The thread formation of a silk protein is usually governed by complex rheological viscoelasticity in the spinning duct. Prior to fiber spinning the fluid silk needs to be conveyed from the ampullate sac into a tiny but lengthy spinning duct (Vollrath and Knight, 1999). The main issue is the flow resistance caused by the convergent flow of fluid silk around the funnel. The approximate pressure required to push liquid silk over the spinning duct could be estimated by the Hagen-Poiseuille equation (Bird et al., 2002).:

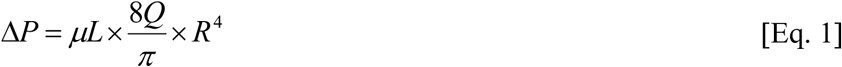

where Δ*P* is pressure drop, *L* is a length of tube, *R* half diameter and *Q* is flow rate.Eq. 1 shows a 4^th^-order influence of vessel radius on flow resistance for a Newtonian fluid within a tubular configuration. The diameter of the spinning duct shows greater than tenfold reduction than that of the sac-like mid-piece, so that the flow resistance in the duct could increase drastically, up to 10,000 times. Similar findings related with the high flow resistance in spiders were also reported by Kojic et al. (Kojic et al., 2006). The sac does not, however, exhibit any musculature tissue to facilitate expelling the secretions (personal observations) for compensating the huge extrusion resistance. Additionally, water removal in the spinning duct that further increases the concentration and viscosity of silk dope could even unfavorably magnify the obstacle of spinning. In summary, according to these arguments, there might exist alternative physiological pathways that dissipate the silking resistance built up along the spinning duct as well as promote the overall spinning efficiency. Therefore, a reevaluation of the spinning process of spider silk threads is highly desirable.

In this study, we assessed the liquid/solid concentration of silk along the spinning system, silk entrance at gland sac vs. silk exit at the spinneret, as a clue for the overall liquid flow and silk physiology of the spider. This, in turn, will provide a beneficial means to elucidate the global pinning mechanism of spider silk proteins; therefore, also shed light on the fabrication of future silking process.

## Materials and Methods

### Spider silk collection and mechanical properties measurement

Adult female *Nephila pilipes* (Fabricius 1793) (body length > 40 mm) were collected from the low-altitude mountainous areas in northern and central Taiwan. In this study, we first examined the basic physical properties of their silks. Only dragline silks produced by major ampullate glands were utilized. To collect the dragline silk, we pulled the silk from the spinnerets of a secured spider under a dissecting microscope then taped the two threads on a rotor powered by a motor (Blamires et al., 2013). A digital winder controlled the take-up speed, ranging from 1.6 – 8.0 m/min. The fineness of silk fibers was measured and expressed in denier (d) defined as the weight (in gram) of a fiber with a length of 9000 meters. The silk fiber tenacity was obtained with a tensile tester (ZWICK 1445, Zwick Roell). A gauge length of 25.4 mm with a crosshead speed of 100 mm/min was utilized. In each measurement, 20 silk filaments were used for the tensile measurements and the all of the results were subsequently averaged to obtain the representative data.

### Liquid content measurement of silk fibers exiting the spinneret

Each *Nephila pilipes* spider was first secured on a cardboard, placed on an analytical balance (SX205 Delta Range, Mettler Toledo) with readability of 0.01mg. The balance was connected to a personal computer to collect the real-time weight change of the spider during the experiment. Prior to silking, weight loss of the spider was recorded for 30-60 minutes at resting condition, and the metabolic weight change was acquired. The major ampullate gland silk threads of the same spider were then harvested using the reeling rotor for 30-90 minutes; *in situ* and real-time weight change of the same spider during forced-silking was traced, too. Subsequently, the collected silk threads were further dehydrated in the oven and the dry weight was quantified. Finally, the liquid content of silk immediately extruded from the spinneret can be calculated by the following equation:

7

Liquid content 
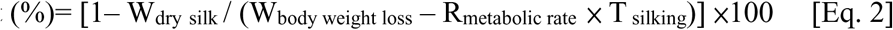

where Wdry silk designates the dry weight of silk drawn from the spider, W_dry silk_ designates the body weight loss of a spider during the forced-silking process, R_metabolic rate_ designates the average weight loss rate of the spider due to metabolism, and T _silking_ designates the time period of silking. Total five biological replicates were performed with separate spiders, followed by the liquid content estimation respectively.

### Measurement of gland silk liquid content

The liquid content of the gland silk was estimated by a thermogravimetric analyzer (TGA-Q50, TA Instrument Co., USA). Major ampullate glands were dissected directly from female *N. pilipes* under the dissecting microscope and the procured glands were gently-wiped with Kimwipes to remove residual fluid outside the glands. The shell of the silk gland was carefully opened and then removed with tweezers. The weight loss of fluid silk during heating was monitored and the heating temperature ranged from 25 ∼ 600 °C. During the initial stage of continuous heating, the evaporation of liquid resulted in a plateau of mass loss value and thus the ratio of dehydrated silk to liquid was estimated.

## Results

Major ampullate dragline silks of *N. pilipes* were collected at varied winding speeds via a programmable reeling rotor. Physical properties of the silk samples, both fiber thickness and tensile strength, were systematically assessed (Fig. 1). Fig. 1A and 1B indicate the take-up speed dependence of silk fiber tenacity and diameter, respectively. The average silk fiber tenacity of *N. pilipes* was around 7.6 – 8.4 g/denier and the utmost value was between 10 – 12 g/denier. No evident speed-tenacity relationship was observed in Fig. 1A. On the contrary, the thickness of silk fibers drawn from the spiders seemed to be influenced by the winding speed (Fig, 1B)(Davies et al., 2013). At low-take up speed (< 3m/min), the diameter of silk threads slightly increased along with the spinning speed; the diameter of spun fiber gradually declined with the spinning speed when it exceeded 3 m/min (e.g. ∼1.1 denier at 3 m/min vs. 0.7 denier at 8 m/min).

**Figure 1.**
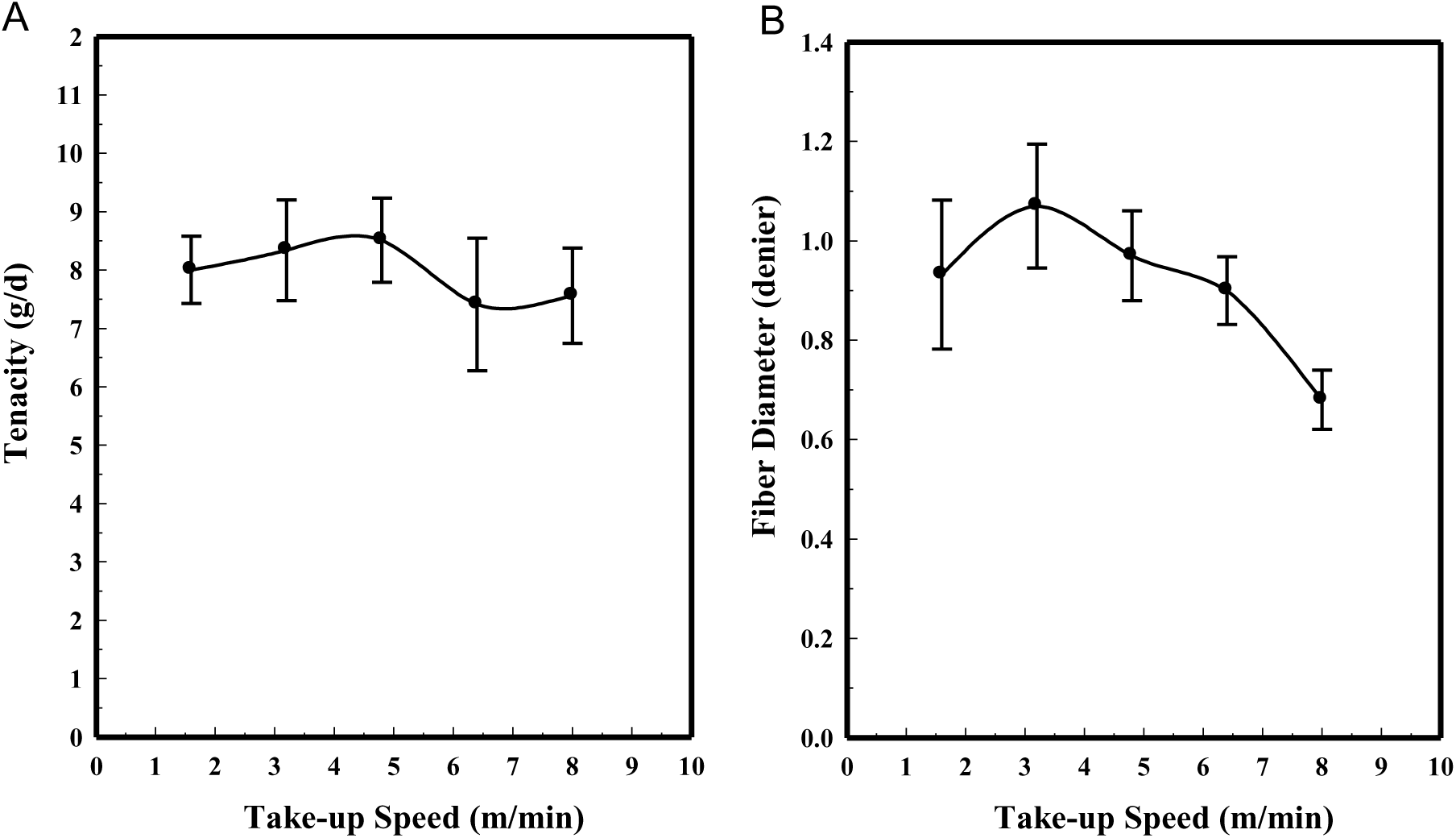
Physical properties of dragline silk from *N. pilipes.* (A) Tenacity (g/denier) and (B) diameter (denier) under different take-up speeds.

To gain more physiological insights on the silk spinning process of a spider, we measured the concentration change of silk dope along the spinning process. In Fig. 2, two approaches were utilized to gauge the concentration at both terminals of the silk gland system: the gland sac (as silk source) and silk fibers leaving the spinneret (as silk drain). Briefly, in Fig. 2A left, a major ampullate silk gland was isolated from the abdomen of a dissected *N. pilipes*. Similar to other *Nephila* spider, the ampullate silk gland of *N. pilipes* also consisted of a tail, a sac-like mid-piece and a thin looped spinning duct, which links the sac to the funnel. We then further analyzed the solid/liquid content of amupllate sac portion via TGA and the result was shown in Fig. 2B. During heating, the sample mass loss reached the first plateau (50.0 wt%) when the temperature exceeded 100℃. The level-off of mass during this stage of heating was most likely as a result from liquid evaporation. When the heating temperature exceeded 300℃, thermal degradation occurred.

**Figure 2.**
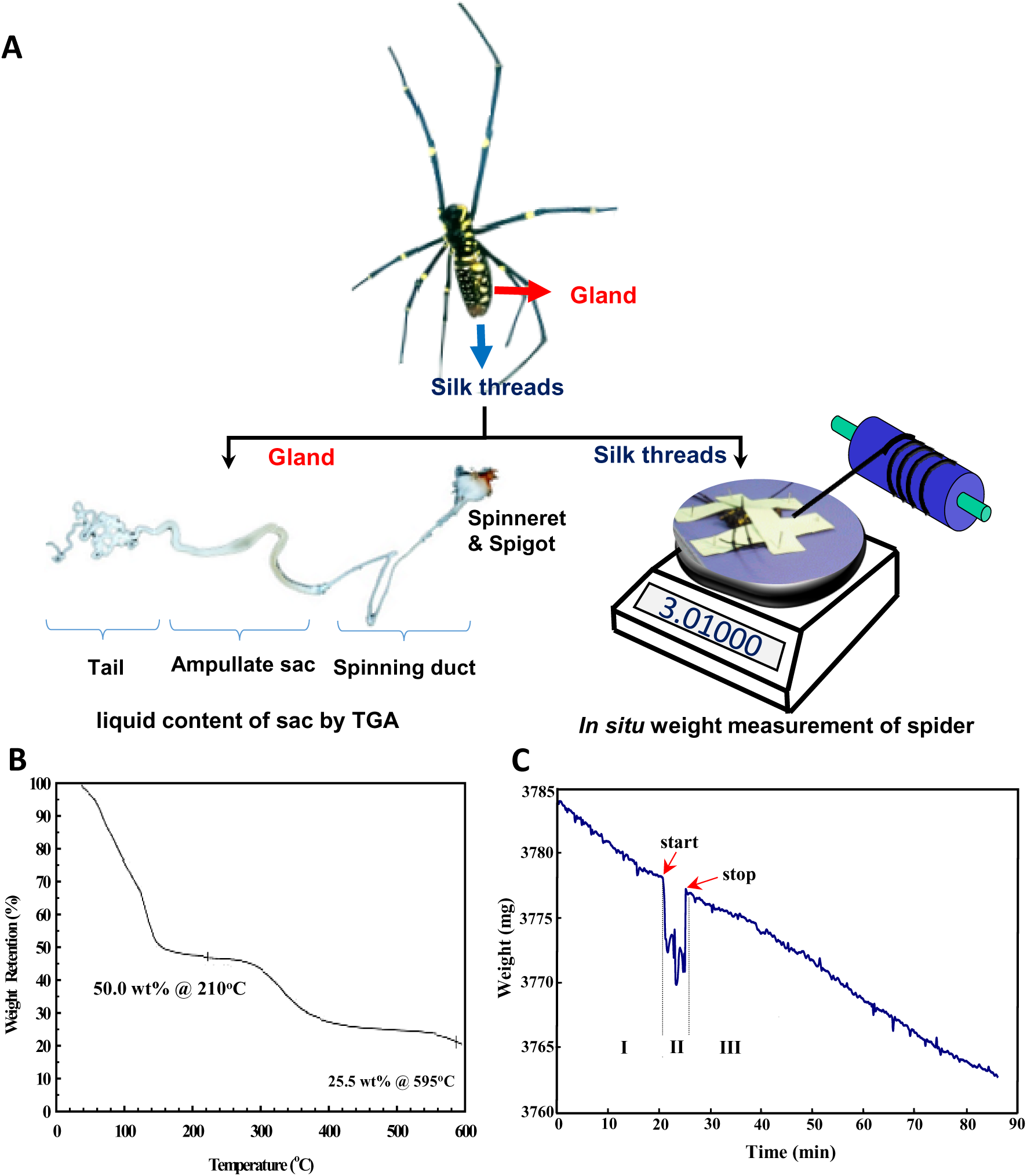
Measurements of spider silk concentrations of *N. pilipes* along the spinning process. (A) Schematic for assessing liquid concentration of the silk dope at different locations of the spinning pathways. The concentration of silk at the gland sac harvested from a spider was estimated via TGA (left). The concentration of silk exiting the spigot was gauged by monitoring the weight loss of the spider on the microbalance and the collected dried silk (right). (B) TGA weight percentage of the liquid silk sample indicated from Fig. 2A left. (C) Real-time body mass change of the spider in the process of (I) pre-silking, (II) peri-silking and (III) post-silking upon forced-spinning via the winder.

Depicted in Fig. 2A right, a novel strategy was implemented for an accurate evaluation of the liquid content of silk exiting the spigot of the spider’s abdomen. The quantity of liquid carried by the silk fibers exiting the body was indexed by subtracting the amount of dried silk from the spider weight change associated with the silking process. Specifically, the spider was securely immobilized onto the microbalance with the major ampullate silk fibers attached to the reeling winder. The body weight of the spider was then continuously monitored *in situ*. As an example demonstrated in Fig. 2C, the change of body weight of the spider was recorded from before to after the forced-silking process. The real-time weight loss curve was divided into three states: state I, II and III which indicated the pre-, peri‐ and post‐ silking process respectively. In states of I and III, the secured spider was kept statically on the microbalance without experiencing the dragging force exerted by the winder. The weight decreased monotonically at the average rate of ∼0.3 mg/min, the respiratory metabolic rate of the spider (Anderson, 1970; Kawamoto et al., 2011). In state II (the silking period), the dragline silk was drawn by the controlled winder and the reading of the microbalance was abruptly reduced, owing to the disturbance of the external force. Nevertheless, the mass reduction of *N. pilipes* was obtained by estimating the weight loss of state II, from the start to the end of silking. The silk fibers collected from the winder (Fig. 2A right) were harvested and dried in an oven; the solid content of those desiccated fibers was gauged subsequently. Finally, the actual liquid amount associated with silk fibers was estimated by Eq. 2 which represents a quantitative means for probing the percentage liquid content of silk fibers *in situ*.

Table 1 summarized the detailed description of both liquid and solid content of silk fibers from *N. pilipes* spiders. Five replicate measurements were independently conducted with individual spiders. The estimated metabolic rates were 0.3∼0.5 mg/min and the liquid contents of dragline silk leaving the spinneret ranged from 80.9 to 96.1 %. This, in turn, suggests an overall higher liquid content of silk dope at the spinneret than that of silk at the gland sac, ∼50.0 wt%.

**Table 1.** Summary of liquid and solid content estimation on major ampullate dragline silk immediately drawn from the spigots of female *N. pilipes* spiders.

| **Item \ Spider** | **# 1** | **# 2** | **# 3** | **# 4** | **# 5** |
| --- | --- | --- | --- | --- | --- |
| **Weight of spider (mg)** | **3041.0** | **2850.6** | **3013.6** | **2698.1** | **2873.1** |
| **Weight loss during silking (mg)** | **12.0** | **15.5** | **44.8** | **33.1** | **12.6** |
| **Metabolic rate (mg/min)** | **0.3** | **0.3** | **0.4** | **0.5** | **0.3** |
| **Silking time (min)** | **29.0** | **25.0** | **31.0** | **23.0** | **30.0** |
| **Weight of dried silk (mg)** | **0.5** | **0.7** | **1.3** | **1.2** | **0.6** |
| **Silk conc. exiting spinneret (wt%)** | **19.1** | **7.4** | **3.9** | **5.4** | **12.0** |
| **Water conc. exiting spinneret (wt%)** | **80.9** | **92.6** | **96.1** | **94.6** | **88.0** |

## Discussion

Up to 46,856 species of spiders have been identified to produce silks (World_Spider_Catalog, 2017).In this study, we implemented a series of spider screening and picked golden orb-weaver *N. pilipes* as our model species: abundance in population, ease of handling and possession of silk with strong mechanical properties (Blamires et al., 2012). *N. pilipes* is among the largest orb-web builders worldwide, especially in Taiwan, whose body length is in the range of 30-50 mm. The maximum amount of silk from a healthy adult can sometimes reach 5-15 mg by a single harvest using forced-silking approach (Tso et al., 2005). The average silk fiber tenacity of *N. pilipes* (7.6 – 8.4 g/denier) is comparable to that of *N. clavipes*’s dragline silk (Blamires et al., 2012; Vollrath and Knight, 2001a).

Controlling dragline fiber thickness of a spider has been considered involving in a complex regulating machinery within its spinneret. Our forced-reeling approach provided a means to further reveal the regulation. From Fig. 1, we speculate such thickness pattern may result from the internal regulation of the spinneret to release the pressure caused by forced-silking on the spiders. Vollrath and Knight (Vollrath and Knight, 2001a) suggested a valve system at the exit of the spinneret is regulated by muscular action and could vary the thickness of the silk thread. Based on the assumption, we further hypothesize that only when the take–up speed exceeds the threshold beyond the spider’s internal regulation capability (eg. > 3m/min), the diameter of spun fiber might be reduced with spinning speed. In nature, a spider could easily apply high spinning speed by administering sufficient external shear forces, dragging through its own leg or descending via gravity, to produce the silk threads of need (Davies et al., 2013).

The emphasis of this research was mainly to elucidate the physiological regulation of silk concentration along the spinning apparatus of a spider. In the perspective of material mass balance, we regarded the silk gland as an open system and probed silk and liquid concentrations at both ends. The liquid content of silk dope inside the ampullate sac (designated as the system inlet) was estimated to be ∼50.0 % (Fig. 2B), similar to that of *N. clavipes* (Hijirida et al., 1996); on the other side, the liquid content of the silk threads exiting the spigot (designated as the system outlet) was calculated 80.9-96.1% (Table 1). As noted in Eq. 2, the devised estimation, we believe, is able to approach the genuine liquid content of spider silk right at the exit of the spinneret upon spinning. The calculation is imply based on the material balance of liquid in respect of two interplayed elements: silks and spiders. Namely, the total weight loss of the spider during silking equals to the summation of silk, liquid and spider metabolism. Therefore, the liquid portion can be accurately assessed by considering those factors.

Demonstrated again in Table 1, our study focused on evaluating the changes of liquid content of silk immediately after it was drawn from the spinneret, and thus avoided the biased measurement due to uncontrolled liquid evaporation from the silk fibers over time. We propose that the majority of liquid co-spun with the silk threads could be quickly lost once the moisture is exposed to the ambient environment, due to the high surface-to-volume ratio of the dragline silk (microns of thickness). Liquid removal by evaporation only takes place after the silks exit the spinneret that can serve as an important mechanism for facilitating silk thread formation.

Furthermore, the increase of liquid content of the silk dope along the spinning process, ∼ 50.0% at ampulla sac vs. 80.9-96.1% at the spigot, implies the overall liquid supplementation mechanism might be involved in the spinning process. In complementary to the liquid extraction model at the spinning duct of a spider, we speculate there is a liquid addition mechanism at the distal portion of the spinning duct, adjacent to the spigot part (Vollrath and Knight, 2001b). The extra liquid supplying mechanism, in turn, could possibly abrogate the huge silking resistance of silk dope resulting from the tapered spinning duct as well as the hardened silk threads upon silking. Besides, the added liquid layer exerted on the spider silk threads could also serve as a protective coating to alleviate the potential damage of lining layer of the spinning duct upon silking. As schemed in Fig. 3, a highly facile spinning process is proposed in two steps: (1) liquid silk dope stored in the ampulla sac is first drawn through the spinning duct where liquid-to-solid silk transition is expedited by water removal and ion exchange; and (2) liquid supplementing at spigot to reduce the silking resistance and further facilitate the silk threads extrusion.

**Figure 3.**
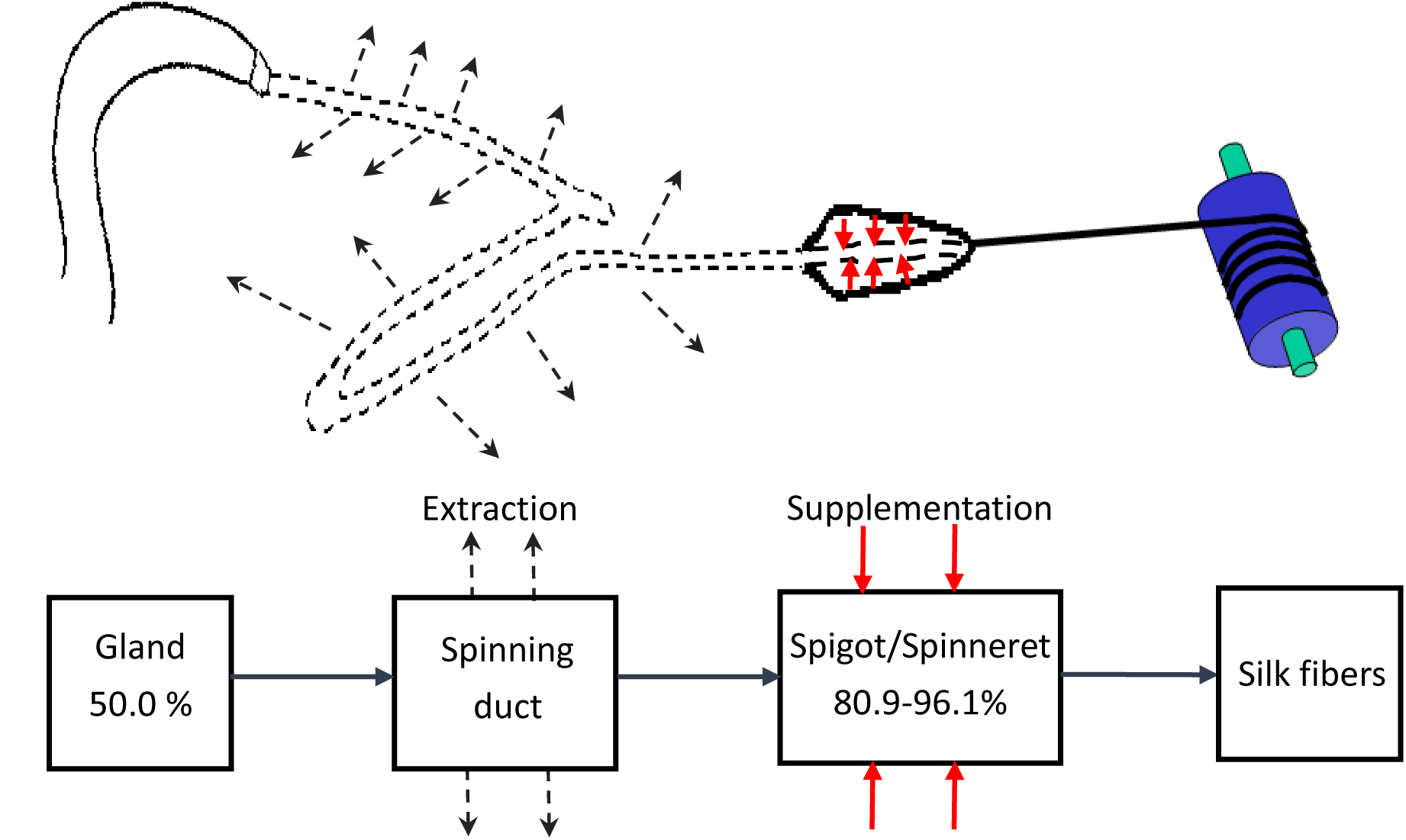
Schematic of the proposed major ampullate silk spinning model of *N. pilipes.* The liquid content of the silk dope decreases initially as it passes through the spinning duct (liquid extraction; dashed line), and then increases again at the spigot (liquid supplementation; red line) upon the silk threads formation.

To sum up, we propose an alternative working hypothesis for the spigots of a spider that can macroscopically supplement liquid into the silk dope as it is transported from the sac to the spinneret. The novel finding presented here might provide more understandings of the spider silk physiology as well as alternative insights to advance the current spinning technologies for the man-made spider silks.

## Acknowledgements

The authors would like to acknowledge the finical support of Ministry of Science and Technology of Taiwan (MOST 104-2221-E-038-003P; MOST 104WFA0163944).

